# Neural signatures of harm aversion predict later willingness to exert effort for others’ rewards

**DOI:** 10.1101/2025.10.03.680267

**Authors:** Luis Sebastian Contreras-Huerta, Hongbo Yu, Annayah M. B. Prosser, Patricia L. Lockwood, Felipe Rojas-Thomas, Molly J. Crockett, Matthew A.J. Apps

**Affiliations:** Center for Social and Cognitive Neuroscience (CSCN), School of Psychology, Universidad Adolfo Ibáñez, Viña del Mar, Chile; Department of Experimental Psychology, University of Oxford, Oxford, Oxford OX1 3PH, UK; Centre for Human Brain Health, School of Psychology, University of Birmingham, Birmingham B15 2TT, UK; Center of Social Conflict and Cohesion Studies, Santiago, Chile; Department of Psychological and Brain Sciences, University of California Santa Barbara, Santa Barbara CA 93106, USA; Marketing, Business and Society Division, School of Management, University of Bath, BA2 7AY, UK; Institute for Mental Health, School of Psychology, University of Birmingham, Birmingham B15 2TT, UK; Christ Church, University of Oxford, Oxford OX1 1DP, UK; Department of Psychology, Princeton University, Princeton NJ 08540, USA; University Center for Human Values, Princeton University

**Keywords:** Prosocial effort, harm aversion, anterior insula, temporo-parietal junction, social decision-making, motivation, morality

## Abstract

Prosocial behaviours—actions that incur personal costs to benefit others—are central to human social life. Two key domains are moral harm aversion, where individuals forgo personal gains to prevent harming others, and prosocial effort, which involves exerting effort to benefit others. Although previous studies suggest a relationship between these behaviours, it remains unclear whether neural responses in one domain can predict prosocial motivation in another. Here, we tested whether neural sensitivity to morally salient information in harm aversion could predict prosocial effort later. Participants completed two tasks: a harm aversion task during fMRI, in which they traded off monetary profit against delivering electric shocks to another person; and, one week later, a prosocial effort task outside the scanner, in which they decided whether rewards for others were worth the required physical effort. We focused on three regions implicated in cost–benefit decision-making and social cognition: the anterior cingulate cortex (ACC), anterior insula (AI), and temporoparietal junction (TPJ). Behaviourally, greater harm aversion was associated with increased prosocial effort. Neurally, AI responses to others’ harm predicted sensitivity to others’ rewards in the effort task, consistent with a role in representing others’ outcomes across positive and negative valences. By contrast, TPJ responses to profit from harming others predicted decreased sensitivity to others’ rewards, suggesting a role in context-dependent valuation that may constrain prosocial behaviour. These findings demonstrate that neural responses to morally salient information in one context correlate with prosocial motivation in another, highlighting mechanisms that bridge moral sensitivity and effortful prosociality.

## INTRODUCTION

Prosocial behaviour—costly actions that benefit others—is fundamental to human social life^1^. It promotes cohesion and cooperation, facilitating the functioning of communities and societies^2,3^. Yet, individuals vary in their willingness to act prosocially, depending not only on the personal cost but also on the nature of the benefit provided to others. For example, in a moral context, people are often willing to forgo personal gains to prevent others’ harm. However, they might show less willingness to exert effort to provide rewards for others in less normative charged contexts^4,5^. Still, some individuals consistently act prosocially in both moral and effort-based contexts^6^. This suggests that sensitivity to morally-relevant information could also relate to prosocial motivation in other domains. What kinds of morally salient signals in one context might forecast prosocial tendencies in others?

A central source of moral salience is the potential harm that others might experience. From early childhood, people learn the moral norm of “do no harm,” a proscription that is strongly internalised across different cultures^7–11^. Avoiding harm to others is often prioritised over self-interest, leading individuals to sacrifice personal gains to prevent others’ suffering^12,13^. Indeed, decisions that involve causing harm—even hypothetically— are typically experienced as aversive, morally salient, and norm-violating^14–17^. Consistently, Crockett and colleagues^4,18,19^ found that individuals were willing to forgo more money to avoid inflicting pain on others than on themselves—a phenomenon termed “hyperaltruism.” These findings suggest that people are highly reluctant to harming others, even at a personal cost.

Unlike harm aversion, however, working to benefit others through effort is not typically guided by a moral norm. Effort is inherently costly, and effort-based decisions generally follow cost-benefit principles: as the required effort increases, so must the anticipated reward to justify the action^20–22^. When the rewards accrue to another person, rather than oneself, the cost–benefit trade-off becomes even steeper^5,23,24^. Supporting this, Lockwood and colleagues^5^ showed that, although people will exert effort to benefit others, they are more willing to work for their own rewards than for others’, a tendency termed “prosocial apathy.” Thus, whereas people are often motivated to forgo personal gains to prevent others’ harm, they are less motivated to incur costs when no moral wrongdoing or potential for harm is at stake. Indeed, Volz and colleagues^25^ found that willingness to harm oneself to reward others was weaker than to reward oneself, and uncorrelated with harm aversion. These findings suggest that prosocial motivation depends on the context: people are generally more motivated to prevent others’ suffering than to effortfully promote their wellbeing. This dissociation might rely on distinct mechanisms. In harm aversion, people act on a moral imperative to avoid harm, where profiting at the expense of others’ suffering may diminish the subjective value of personal gains^19,26,27^. By contrast, prosocial effort depends on valuing others’ rewards enough to outweigh effort costs, as individuals typically prioritise self-benefitting actions^28^. However, could sensitivity to morally salient information in one domain—such as harm aversion—predict motivational tendencies in another, such as prosocial effort?

Prior research suggests that even though harm aversion and prosocial effort may be driven by different motives, they are supported by distinct, but potentially overlapping, psychological and neural mechanisms^6,7,29,30^. A recent online behavioural study^6^ found that individuals who were more hypothetically averse to harming others also tended to be more willing to exert effort to help others, with shared variance best explained by traits linked to empathic concern and affective sensitivity. These findings suggest that an other-regarding mechanism—based on sensitivity to others’ outcomes regardless of valence— may generalise from a moral to a reward-seeking context. From this perspective, individuals who are more sensitive to the moral salience of another’s potential harm may also be more sensitive to others’ rewards, perceiving them as more worth the effort.

Converging evidence points to the anterior cingulate cortex (ACC), anterior insula (AI), and temporoparietal junction (TPJ) as key candidates for this other-regarding mechanism—these regions are engaged during vicarious pain, when tracking others’ rewards, and when inferring about others’ mental states^31–45^. Crucially, individual differences in their activity predict variability in prosocial choices, including harm aversion, effort-based and reward-delivery contexts^24,40,41,46–54^. Thus, these areas may encode salient affective signals, such as others’ harm or rewards, that motivate behaviour^31,35,38,55–61^. From this perspective, neural sensitivity to others’ outcomes— whether positive or negative—may guide prosocial decisions. For example, responsiveness to others’ harm in a harm-aversion context could serve as a trait-like marker of moral sensitivity and predict how motivated people are to seek rewards for others. Yet, it remains unclear whether neural responses to one outcome valence (e.g., others’ harm in a moral context) can predict prosocial tendencies toward another (e.g., exerting effort for others’ rewards in a reward-seeking context).

Another source of moral information arises when individuals gain benefits through immoral means—for example, earning profit by causing others to suffer. Here, the focus is not on the harm itself but on the profit derived from ill-gotten means, which can serve as a marker of one’s moral principles. From this perspective, an alternative interpretation proposes that the ACC, AI, and TPJ may not exclusively encode others’ outcomes, but instead participate in a broader valuation system that integrates profit with moral context^15,41,62–68^. These regions, which are engaged in domain-general value coding of rewards, are sensitive to the context, particularly in social settings^24,41,47,50,54,68–72^. Crucially, when personal profit is tied to harming others, the subjective value of self-reward may be attenuated in these regions, reflecting the moral cost of such gains. hus, individuals who retain strong neural responses to profit despite its harmful consequences may experience less moral conflict—potentially predicting lower prosocial motivation in other domains.

Supporting this view, Crockett et al.^19^ showed that participants with reduced striatal responses to profits from harming others were more hyperaltruistic, suggesting that moral transgressions can diminish the subjective value of personal gain. Although the ACC, AI, and TPJ were also recruited during harmful decisions in that study, their activity did not directly predict behaviour. Nonetheless, responses in these regions may generalise to situations where moral conflict is less explicit, such as deciding whether to exert effort to benefit others. However, to our knowledge, no study has yet tested whether neural activity during harm aversion predicts prosocial effort. Demonstrating such a relationship would indicate that responses in these areas—whether reflecting moral-conflict sensitivity or an other-regarding mechanism—constitute a trait-like signature that extends beyond moral contexts. This would support the idea that valuation processes in these regions bridge different social decision-making domains to shape prosocial behaviour.

In this study, we tested whether moral harm aversion and prosocial effort are linked at both behavioural and neural levels. Specifically, we asked whether individual differences in neural responses during harm aversion could predict subsequent motivation to exert effort for others. Participants first completed a harm aversion task during fMRI, choosing whether to profit at the expense of painful shocks delivered to themselves or another person. One week later, they performed a prosocial effort task, deciding whether to exert varying levels of physical effort to obtain rewards for themselves or others. We focused on neural responses in the ACC, AI, and TPJ to others’ harm and to profit from harming others, and examined whether these signals predicted the degree to which participants valued others’ rewards in the effort task. By linking neural sensitivity to morally salient outcomes during harm aversion with subsequent prosocial motivation in a different domain, this study highlights how specific moral neural signatures may forecast prosocial behaviour across contexts, advancing our understanding of the motivational mechanisms that support human prosociality.

## METHODS

### Participants

80 healthy volunteers (aged 18–38 years) were recruited from the University of Oxford and the local Oxford community. The study was approved by the University of Oxford ethics committee (R50262/RE001). Participants were excluded if they had a history of neurological or neuropsychiatric disorders, psychoactive medication or drug use, or were pregnant. Additionally, individuals who had previously participated in studies involving social interaction or had studied psychology for at least one year were excluded to minimise potential biases in psychological and neural processes. All participants provided written informed consent and were compensated for their time (£15 per hour for the fMRI session and £8 per hour for the behavioural session), along with additional bonuses earned through task decisions.

From the initial sample, 66 participants (mean age = 22.6, SD = 3.7; 38 females) were included in the final analysis. Four participants did not attend the behavioural session. Two were excluded for expressing doubts about the social manipulation and the existence of the receiver. Seven were removed due to excessive head motion during scanning or technical issues with the harm aversion task. One additional participant’s neuroimaging data was lost due to a scanner malfunction.

### General Procedure

Participants completed a multi-stage study investigating social behaviour and cognition. As part of this study, they performed the harm aversion task during an fMRI scanning session and, at least one week later, the prosocial effort task in a separate behavioural session. Some subsets of these data have previously been reported in Yu et al.^73^, Contreras-Huerta et al.^74^, and Zho et al. (*in press*). However, the present manuscript addresses a distinct research question, and none of the neuroimaging or behavioural analyses reported here have been included in those publications.

#### fMRI session: Harm aversion task

Prior to the fMRI session, participants completed a battery of online personality questionnaires. At least one week later, they attended the MRI scanning session. At the beginning of the session, participants underwent a pain thresholding procedure designed to (i) familiarise them with the level of pain to be traded off in the task and (ii) identify a pain intensity that was subjectively comparable across participants^4^. After this procedure, a role assignment protocol was applied, where participants were randomly assigned the role of the Decider and another unknown person, who was actually a confederate, was assigned the role of the Receiver^4,5^ (see **Supplementary Methods**). Participants were instructed that their decisions would have consequences for both themselves and the Receiver, and that all choices were confidential and anonymous to minimise concerns about reputation or reciprocity. They were also informed that receivers would complete a separate set of tasks that did not involve outcomes for other people.

After practicing six trials, participants completed the harm aversion task while in the scanner **(Fig. 1A).** In this task, participants traded-off units of pain, represented by a number of electric shocks, against profit, in the form of money. In each trial, participants made a series of binary choices where one option, the *harmful* option, contained more electric shocks in exchange of higher amounts of money, while the other option, the *helpful* option, was associated with fewer shocks and less money. The money was always received by the participants themselves, i.e. the Deciders, but the shocks were allocated to an unknown person on half of the trials, the Receiver (i.e. *other* condition), and to the participant in the other half of the trials (i.e. *self* condition). Importantly, no shocks nor money were delivered inside the scanner. Instead, one randomly selected trial was implemented outside the scanner. If the selected trial was a self trial, participants received the corresponding electric shocks. If it was an other trial, participants were told that the Receiver would receive the shocks in another room—though, in reality, no shocks were administered, as the Receiver was a confederate. The monetary reward from the selected trial was added to participants’ total earnings and given to them at the end of the behavioural session.

**Figure 1.**
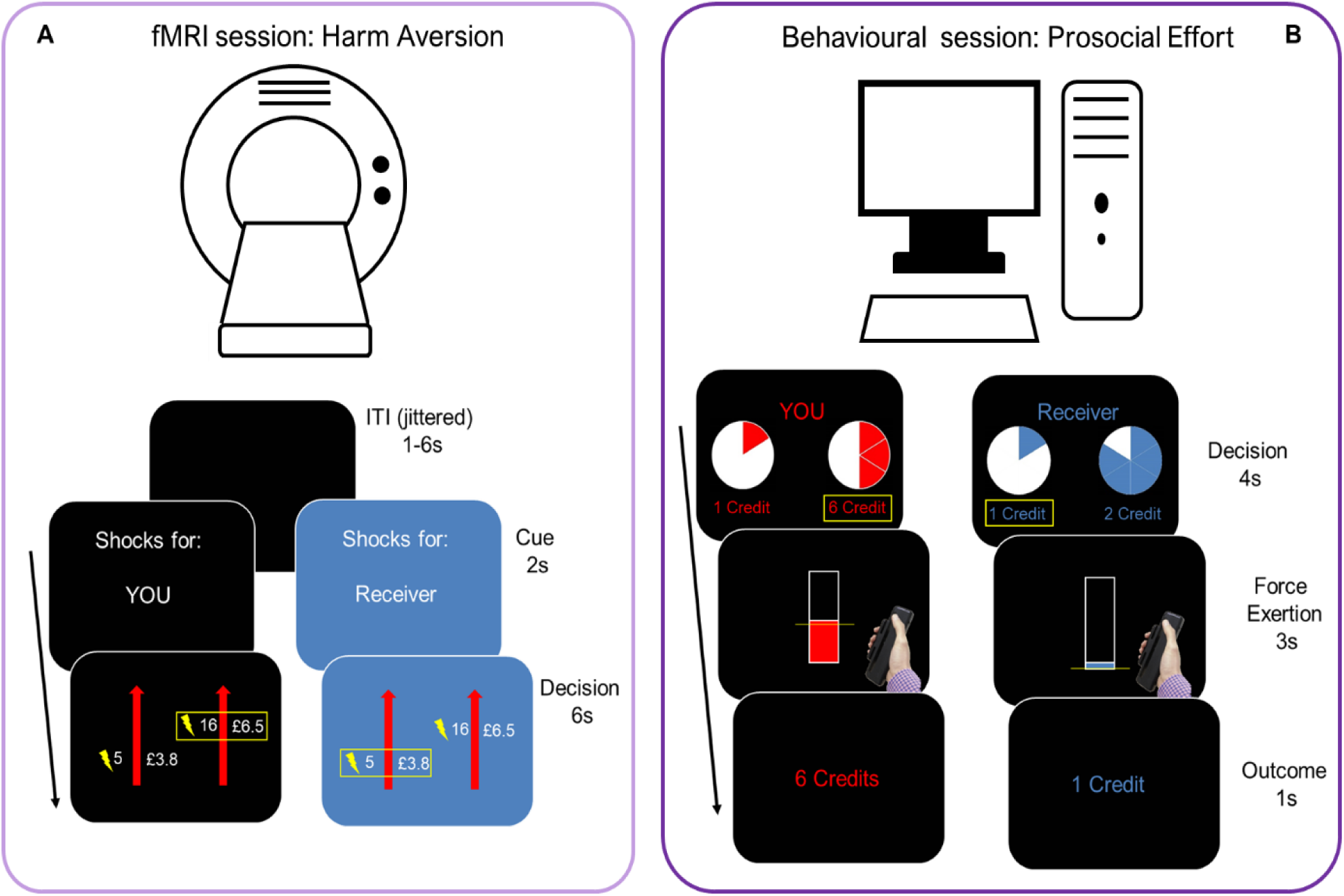
General Procedure. *A. fMRI session - Harm Aversion Task*. Participants completed the harm aversion task while in the scanner, where they decided between a ‘helpful’ option, associated with a lower amount of money and shocks, and a ‘harmful’ option, with more money for more of shocks. In half of the trials, shocks were delivered to the participants themselves (*self* trials, black background), while in the other half, shocks were intended for the Receiver (*other* trials, blue background*). B. Behavioural session – Prosocial Effort.* At least one week later, participants completed a behavioural session where they made prosocial effort decisions. In each trial, they chose between a ‘rest’ option (0% of their maximum voluntary contraction - MVC), associated to 1 credit, and a ‘work’ offer, with variable higher effort (30-70% of their MVC) and higher reward (2- 10 credits) associated to it. If they chose the work option, they had to exert the required force for at least 1 second within a 3-second window to earn the reward; failure to do so resulted in no reward. In half of the trials, participants received the reward themselves (*self* trials, red background), while in the other half, the reward was allocated to the Receiver (*other* trials, blue background).

The procedure for generating choice options is detailed in Crockett et al.^4,19^. Trials were optimised to efficiently estimate harm aversion parameters (*κ*, see below) while ensuring that profit and pain were uncorrelated (|r| < 0.07, p > 0.5) across trials following the criteria given in Crockett et al.^4^. Four different sets of 72 trials were created and counterbalanced across participants. Each set included four catch trials, in which the harmful option offered less money than the helpful option, resulting in 76 trials per set. In half of the trials, the harmful option appeared on the right side of the screen, and in the other half, it appeared on the left. Finally, all 76 trials were duplicated for both self and other conditions, yielding a total of 152 trials, presented in two runs of 76 trials each (half self, half other per run).

After completing the harm aversion task, participants exited the scanner, and one trial was randomly selected and implemented as described earlier. Participants then completed a debriefing session to assess their understanding and beliefs about the experimental setup. Rather than directly asking whether they doubted the validity of the paradigm, since this could artificially introduce scepticism, we included indirect questions on a 7-point Likert scale (1 = yes, fully, 7 = no, not at all). These questions assessed the clarity of instructions regarding (i) the Receiver’s presence, (ii) the delivery of electric shocks to the Receiver, and (iii) the confidentiality of participants’ decisions. Participants also provided open-ended comments about their experience in the study. Two participants explicitly expressed doubts about the paradigm (e.g., questioning whether the Receiver existed). These participants were excluded from the analysis.

#### Behavioural Session: Prosocial Effort Task

At least one week after the fMRI session, participants attended a behavioural session where they completed several tasks, including the prosocial effort task^5^. They were told that they would continue in their role as Deciders and would be paired with a Receiver from the pool of Receivers in the study. Before making decisions in the main task, participants’ maximum voluntary contraction (MVC) was determined by having them squeeze a handle as strongly as possible. This ensured that effort levels were individually calibrated based on each participant’s grip strength. Afterward, participants experienced each effort level three times to familiarise themselves with the task demands and how effort levels were displayed on the screen.

Next, participants performed the prosocial effort task. In this task, participants traded off different levels of effort against varying magnitudes of financial reward. Monetary rewards were represented as credits, while effort was operationalised as grip force. In every trial, participants decided between two options: a rest baseline, related to no effort and low reward (1 credit); and a work offer, associated with a higher amount of effort that varied across 30-70% of each participant’s MVC, in exchange of higher profit, i.e. 2-10 credits. Crucially, in half of the trials the reward was for participants themselves (*self* trials), while in the other half rewards were for an unknown person, the Receiver (*other* trials). If participants chose the work option, they had to squeeze the handle at the specified effort level for at least one second within a three-second window to receive the reward. Failure to do so resulted in earning zero credits for that trial. If they chose the rest option instead, they simply waited for three seconds and received the baseline reward of one credit. Each unique combination of effort (five levels: 30, 40, 50, 60 and 70% of the participant’s MVC) and reward (five levels: 2,4,6,8 and10 credits) were repeated three times per condition, having 75 trials per beneficiary.

At the end of the session, participants completed debriefing questions similar to those in the fMRI session, assessing their understanding of the experimental setup. They then received reimbursement for their participation, along with any financial bonuses earned throughout the study.

### Analyses of behavioural data

#### Computational Modelling

Computational models were fitted to choices in the harm aversion and the prosocial effort tasks, which precisely quantify the degree in which pain devalues profit and the degree in which money is devalued by effort, respectively. For the harm aversion task, decisions were analysed using a model that calculates the subjective values between harmful and helpful options^4,18,19^. Thus, difference in value between the options can be described as follows:

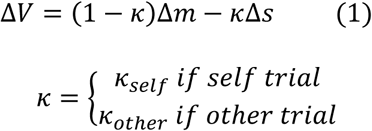

Where *ΔV* is the difference between the subjective value of the harmful and helpful option, and *Δm* and *Δs* are objective differences in money and shocks between harmful and helpful options respectively. In this model, *κ* is a free parameter that represents how harm averse participants are. When κ is 0, participants will accept any offer regardless the pain in exchange. As κ increases and approaches to 1, participants get harm averse, paying more money to avoid additional shocks. When shocks are delivered to the other unknown person, κ_other_ captures the subjective cost of harming the Receiver (other trials), while κ_self_ represents the cost of participants harming themselves (self trials).

For the prosocial effort task, effort-based decisions were fitted into a model^5,23,24^ that measures the degree in which effort discounts rewards parabolically, such that:

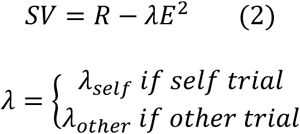

Where *SV* is the subjective value of the work offer with a magnitude of reward *R* and effort level *E*. The free parameter *λ* weights the discount effect of effort on reward, such that when λ is 0, the participant does not discount reward by effort. As λ approaches to 1.5, effort discounts reward in a higher degree. λ was set with a maximum of 1.5 as any value beyond this constraint does not change the predicted decision in the task, i.e. the value of resting (i.e. 1 credit for 0 effort) is higher than the values for the work offer. Finally, the parameter λ represents differently the discount of reward depending on whether it is getting by self (λ_self_) or other (λ_other_).

A softmax function was used in both tasks to transform trial-by-trial differences in value into choice probabilities. For the harm aversion task, the softmax function is used to transform the value of the harmful option relative to the helpful option, ΔV, such that:

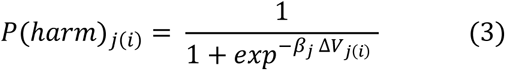

Here, β is a temperature parameter that determines how strongly choices are guided by the value difference (ΔV). Lower β values (approaching 0) indicate greater decision noise, resulting in more gradual, less consistent choice patterns. In contrast, higher β values (approaching 100) produce steeper, near-deterministic choice curves, where the option with the higher value is almost always selected.

For the prosocial effort task, the softmax converts the value of the work offer SV such that:

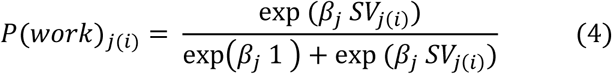

Where *P* is the probability to work of a participant *j* in a trial *i.* When a choice of rest, 1-Pj(i) is used to calculate the probability of not choosing to work.

All parameters described above were estimated individually for each participant using nonlinear optimisation implemented in Matlab (MathWorks) for maximum likelihood estimation.

#### Statistical Analysis

Non-parametric tests were used to compare computational parameters (κ and λ) within and between tasks as these measures were not normally distributed. Computational models were complemented with statistical analyses on raw decision data for each task to separate the effects of different sources of information on choices. Computational parameters represent a specific relationship between costs and benefits in the decision process (e.g., reward being discounted parabolically by effort), integrating both aspects of choice. However, using mixed-effect models (MMs), costs and benefits can be disentangled, being able to test whether the former, the latter, or both, are guiding self and other decisions. Thus, a MM for the prosocial effort task was built, such that:

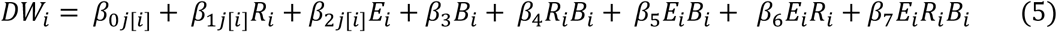

Where decision to work *DW* in the trial *i* is predicted by the fixed effects of reward *R*, effort *E*, beneficiary *B*, and their interactions. Random-intercepts were clustered in subjects *j*, and random slopes on effort and reward were included as it is expected that participants will vary in their sensitivities to this information^5,23,24,54^. Model comparison using AIC between an only random-intercept MM and one with random-slopes confirmed that the latter improves model fitting (random-slopes model = 5830.0, random-intercept only model = 7037.2).

Another MM was built for the harm aversion task, such that:

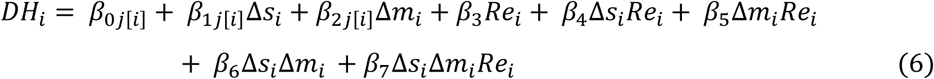

Where decision to help *DH* in the trial *i* is predicted by the objective difference in shocks *Δs* and money *Δm* between options in i, and the recipient of the shocks *Re* for that trial i, together with their interactions. Like the prosocial effort MM, this model had random-intercepts clustered by subjects, and random-slopes for Δm and Δs, as sensitivities to these sources of information are expected to vary across participants^4,18,19^. Looking at the AIC values, model comparison confirmed that adding random-slopes improved model fitting (random-slopes model = 6069.4, random-intercept only model = 7182.0). The outcome variable for all models, i.e. DW and DH, was a binary, factor variable in a logistic MM. These analyses were implemented in R using the *glmer* function.

### fMRI acquisition and preprocessing

Functional MRI scanning was performed on a 3-Tesla Siemens Prisma scanner at the Wellcome Centre for Integrative Neuroimaging (WIN) at The University of Oxford. Functional images were obtained with multiband T2*-weighted echo-planar imaging (EPI) sequence. The EPI images were acquired in an ascending manner, at an oblique angle (approximately 30 °) to the AC-PC plane to minimise the signal dropout in the orbitofrontal areas. The following acquisition parameters were used: 72 slices in interleaved ascending order; matrix size: 108 ×108; voxel size: 2 × 2 × 2 mm^3^ with 1 mm gap; echo time (TE) =30 ms; repetition time (TR) = 1,570 ms; flip angle = 70°; field of view (FOV) = 216 * 216 mm^2^. The structural image was taken using a magnetisation prepared rapid gradient echo (MPRAGE) sequence with 192 slices; TR=1,900 ms; TE = 3.97 ms; field of view = 192 × 192 mm^2^; voxel size = 1 × 1 × 1 mm^3^ resolution. We also acquired a field map (short TE = 4.92 ms; long TE = 7.38 ms; TR = 482.0 ms; resolution = 2 × 2 × 2 mm^3^; FOV = 219 × 219 mm^2^) to correct distortions in the functional images.

MRI data were preprocessed and analysed using SPM12. Functional images were realigned and unwarped with reference to the fieldmap and co-registered to the participant’s own structural image, correcting for distortions caused by susceptibility-induced field inhomogeneities. The EPI images were then realigned and coregistered to the subject’s own anatomical image. The structural image was processed using a unified segmentation procedure combining segmentation, bias correction, and spatial normalisation to the Montreal Neurological Institute (MNI) template; the same normalisation parameters were then used to normalise the EPI images. Finally, images were spatially smoothed with an SPM default Gaussian kernel (8-mm full-width at half-maximum).

### fMRI Analysis

A general linear model (GLM) was constructed to identify brain regions where the BOLD signal varied parametrically with the amount of money and the number of shocks associated with the harmful versus helpful option, for both self and other trials. The BOLD signal was time-locked to the presentation of choice options on the screen. The model included four main regressors, capturing the onset of: (i) *self* trials where the left option was selected, (ii) self trials where the right option was selected, (iii) *other* trials where the left option was selected, and (iv) other trials where the right option was selected. For each regressor, trial duration was defined as the participant’s reaction time (i.e., the interval between option presentation and button press). Each of these four regressors was further modulated by four parametric modulators, reflecting the amount of money and shocks for both the harmful and helpful options, independent of participants’ choices. These parametric modulators were estimated simultaneously, rather than being serially orthogonalized, as is standard in SPM. Regressors of no interest were also included corresponding to onsets of button presses, cue for transitions between conditions, and missing trials, as well as six nuisance regressors to control for head motion. To extract statistical maps for Δs and Δm, we defined contrasts for the first level analysis such that *money*_harmful_ > *money*_helpful_; and *shock*_harmful_ > *shock*_helpful_ for self and other regressors, respectively.

Using the model described above, we extracted statistical maps for self and other conditions from anatomically defined regions of interest (ROIs) using the MarsBar toolbox in SPM. Our analyses focused on five bilateral ROIs: bilateral ACCg, bilateral dorsal and ventral portions of anterior insula (dAI and vAI respectively) and bilateral posterior and anterior portions of temporo-parietal junction (TPJp and TPJa respectively). We used anatomical masks instead of peak voxels for two reasons. First, different portions of these areas can be linked to different processes, and have different connectivity fingerprints^75–77^, making peak voxel comparisons inappropriate. Second, our approach is hypothesis-driven, targeting ROIs that have been previously implicated in cost-benefit decisions, social cognition and prosocial behaviour. Given inter-individual variability, these regions may not exhibit significant group-level effects but could still predict behaviour in another task.

For the AI portions, we used the voxel-wise k-means clustering parcellation by Deen et al.^78^, which divides the insula into three distinct subregions per hemisphere: posterior insula, dAI and vAI. This data-driven parcellation has been independently replicated^79,80^ and widely used in prior studies^54,81,82^. The insula ROIs were acquired in 2mm MNI space from the author’s website (https://bendeen.com/data/). Left and right hemisphere parcels were combined to create bilateral AI masks using the *imcalc* function in SPM12.

For the ACCg, we used thresholded masks from the resting-state connectivity-based parcellation by Neubert et al.^76^. This 21-region frontal cortex atlas was obtained in 2mm MNI space directly from the author website (http://www.rbmars.dds.nl/CBPatlases.htm). The left hemisphere ACCg mask (area 24ab) was mirrored to the right hemisphere to create a bilateral ACCg mask (as the original atlas was derived from participants with a paracingulate sulcus only in the left hemisphere). We excluded posterior voxels from this bilateral mask to better align with the ACCg definition in prior work^75,83^. We focused only on the gyrus portion of the ACC due to its stronger association with social processing, including vicarious pain and social rewards^75^.

Finally, for the TPJ, we used masks from Mars et al. (2012), derived from tractography-based parcellation and resting-state connectivity. The original TPJ parcellation defined posterior (TPJp) and anterior (TPJa) subregions in the right hemisphere. To obtain bilateral TPJ masks, we mirrored the right hemisphere parcels and merged them with their original counterparts. These masks were also obtained from http://www.rbmars.dds.nl/CBPatlases.htm.

Thus, from these five regions (vAI, dAI, ACCg, TPJa and TPJp), we extracted the statistical maps for Δm and Δs in other trials (Δm_other_ and Δs_other_) and correlated them with the behavioural measures of prosocial effort, controlling for multiple comparison across these five regions. We also extracted the statistical maps for self trials (Δm_self_ and Δs_self_) for controlling purposes.

#### Statistical analyses for the association between neural responses and behavioural measures

To test whether neural responses in the harm aversion task predicted prosocial motivation, we correlated beta values from the parametric GLM across five ROIs with behavioural sensitivity to reward in the prosocial effort task, as it has been identified as the main driver for prosocial behaviour^54^. Reward sensitivity estimates were derived from a MM predicting decisions to work for others as a function of reward, effort, and their interaction. This model included random intercepts for participants and random slopes for reward and effort:

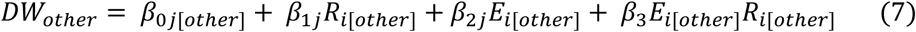

A parallel model using self-benefiting trials was also estimated for control purposes. Neural responses in the five ROIs to money and shocks in the harm aversion task were then correlated with individual reward sensitivity for others (β_1j_R_i[other]_ from the model above). Because several behavioural and neural variables were not normally distributed (Shapiro-Wilk test), we used Spearman partial correlations. To isolate effects specific to other-benefiting motivation, we controlled for reward sensitivity in self trials and corresponding neural responses to self outcomes (Δm_self_ or Δs_self_). Visual inspection of ROI beta distributions revealed a small number of extreme outliers (> ±3 SD). While Spearman correlations are robust to such deviations, we conducted robustness checks excluding these participants. All p-values were corrected for multiple comparisons across ROIs using the false discovery rate (FDR) method^84,85^.

To test whether correlations with fMRI responses aligned with potential behavioural associations between harm aversion and prosocial effort, we conducted an additional analysis. Specifically, behavioural reward beta values for others in the prosocial effort task were correlated with individual sensitivities to money and harm in the harm aversion task. These sensitivity estimates were obtained from a MM predicting decisions to help others as a function of trial-wise differences in money and shocks, and their interaction. This model included random intercepts for participants and random slopes for money and shock differences:

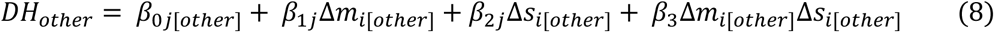

A corresponding model for self trials was also estimated for control purposes. We then tested whether reward sensitivity for others (from the prosocial effort task) was associated with harm aversion–related sensitivities to money and shock in other trials using Spearman partial correlations, controlling for the corresponding self-trial estimates.

## RESULTS

### Hyperaltruism and prosocial apathy effects are correlated

First, we tested whether behavioural results for each task, robustly reported in previous studies, were replicated in the current data set – hyperaltruism, i.e. people being more averse to harm others than themselves for profit^4,18,19^; and prosocial apathy, i.e. people being less willing to put in effort to benefit others compared with themselves^5,23,24^. We took two approaches to test these effects: MMs and computational modelling. For the former, decisions in both tasks were modelled using the different sources of information in each trial as predictors. Thus, for the harm aversion task, trial-by-trial decisions to help were predicted by differences in money between options (Δm), differences in shocks between options (Δs), the recipient of the shocks (self/other), and their interactions (see **Supplementary Table S1** for full results). Using this model, results revealed main effects of Δm (b = -4.22, SEM = 0.4, z = -10.67, p < 0.001) and Δs (b = 1.61, SEM = 0.18, z = 9.17, p < 0.001), having opposite effects on decisions. Thus, greater monetary incentives for harming (higher Δm) reduced helping, whereas greater pain consequences for harming (higher Δs) increased helping. Importantly, there was a main effect of recipient (b = 1.32, SEM = 0.09, z = 15.44, p < 0.001), with participants more willing to help when another person was the recipient of the shocks, replicating the hyperaltruism effect (**Fig. 2A**). Additionally, we found a significant Δm × recipient interaction (b = 0.62, SEM = 0.09, z = 6.86, p < 0.001) indicating that, although higher Δm reduced helping overall, this effect was stronger for self-trials than for other-trials. That is, participants were less likely to harm for high profit when the shocks were delivered to another person (**Fig. 2B**). No other significant effects were found.

**Figure 2.**
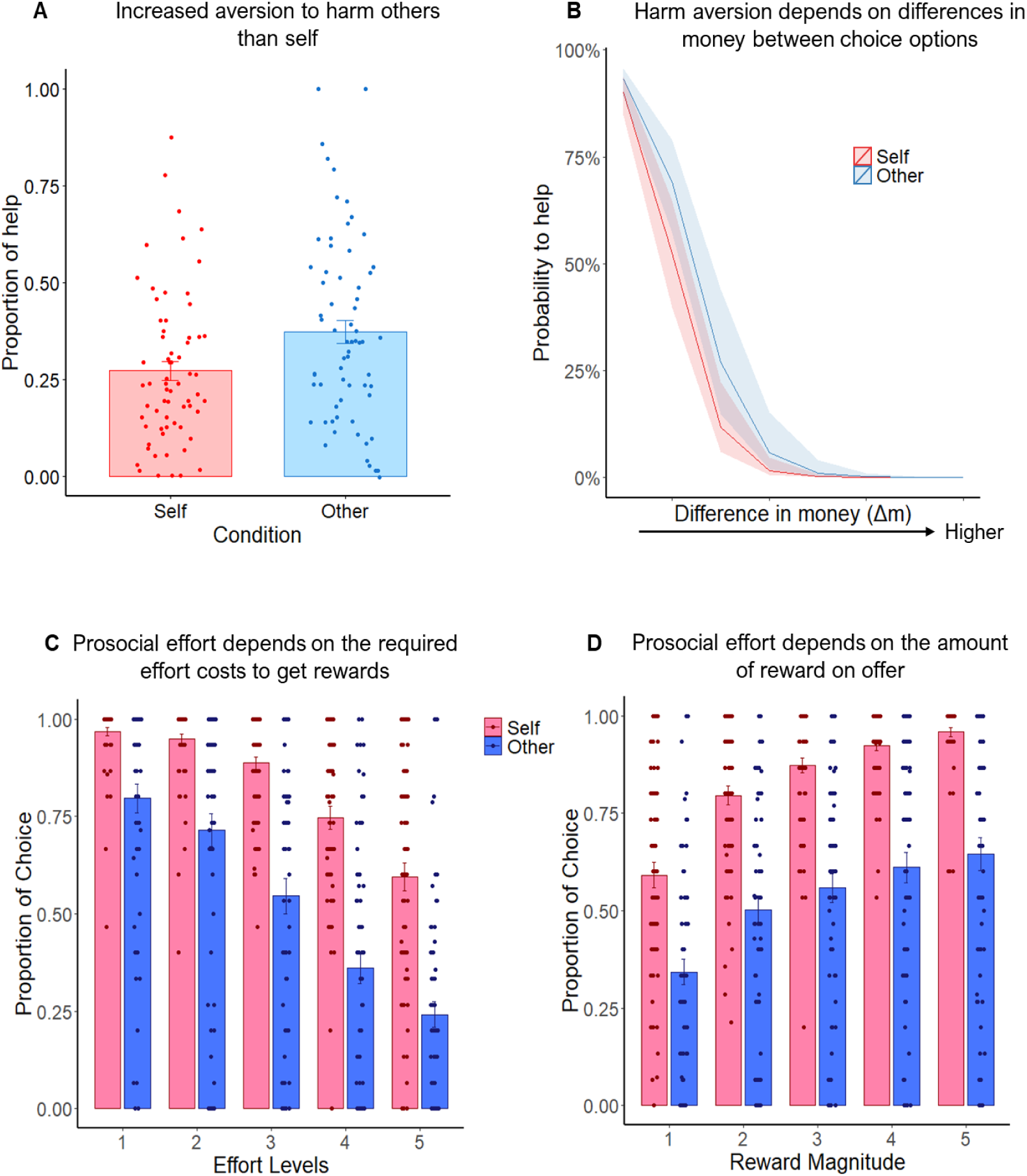
Replication of hyperaltruism and prosocial apathy effects. (A) Participants were more averse to harming others than themselves when trading off electric shocks for profit. The y-axis shows the proportion of helpful choices (i.e., choosing less harm for less profit) relative to harmful choices (more harm for more profit) in the harm aversion task. (B) Model-predicted probability of choosing the helpful option (y-axis) as a function of the monetary difference between options (x-axis). Participants were less averse to harming as the profit for doing so increased, with this effect being stronger in self-trials than in other-trials. Shaded ribbons represent 95% confidence intervals around the predicted values. (C) In the prosocial effort task, participants were less likely to choose the work option at higher effort levels (x-axis), particularly when the beneficiary was another person rather than themselves. The y-axis shows the proportion of work choices (i.e., higher effort for higher reward) relative to the rest option (no effort for a small reward). (D) Participants were more willing to work when the reward was higher (x-axis), especially when the reward was for themselves rather than for others. The y-axis shows the proportion of work choices relative to the rest option. In all plots, dots represent individual participants, and error bars indicate standard errors of the mean (SEM).

For the prosocial effort task, we modelled decisions to work using effort level, reward magnitude, and beneficiary as predictors, along with their interactions (see **Supplementary Table S2** for full results). This analysis revealed main effects of effort (b = -2.52, SEM = 0.21, z = -11.82, p < 0.001), reward (b = 2.22, SEM = 0.18, z = 12.1, p < 0.001), and beneficiary (b = -3.45, SEM = 0.11, z = -32.13, p < 0.001). Participants were less likely to work when the effort requirement was high and when the reward was low. More importantly, interactions between effort × beneficiary (b = 0.26, SEM = 0.1, z = 2.61, p < 0.01) and reward × beneficiary (b = -0.93, SEM = 0.1, z = -9.62, p < 0.001) indicated that these effects were amplified when the reward was for another person (**Fig. 2C, 2D**), replicating the prosocial apathy effect. No other significant effects were found.

Afterwards, we tested whether these effects in trial-by-trial decisions were captured by the parameters obtained from the computational models for the harm aversion and the prosocial effort tasks. First, we compared how averse participants were to harming others relative to self. A Wilcoxon signed-rank test revealed that participants were significantly more averse to harm others (M = 0.38, SEM = 0.03) than themselves (M = 0.27, SEM = 0.03; κ_other_ > κ_self_, W = 504, p < 0.001, **Fig. 3A**), with the majority of participants showing this trend (65.2%, **Supplementary Fig. S1A**). This supported the hyperaltruism effect revealed by the MM. With regards to the prosocial effort task, a similar analysis revealed that participants discounted rewards by effort in a higher degree when the beneficiary was another person (M = 0.28, SEM = 0.04) than themselves (M = 0.06, SEM = 0.007, λ_other_ > λ_self_, W = 19, p < 0.001; **Fig. 3B**). In contrast with the harm aversion task, the vast majority of the participants displayed a self-focused profile, with 95.5% of them being more averse to exerting effort for the other person (**Supplementary Fig. S1B**). Therefore, these results revealed that the same individuals showed a very different behavioural pattern depending on the task-context, i.e. in harm aversion, people are hyperaltruistc, while when they need to be put in effort to work, they are prosocially apathetic.

**Figure 3.**
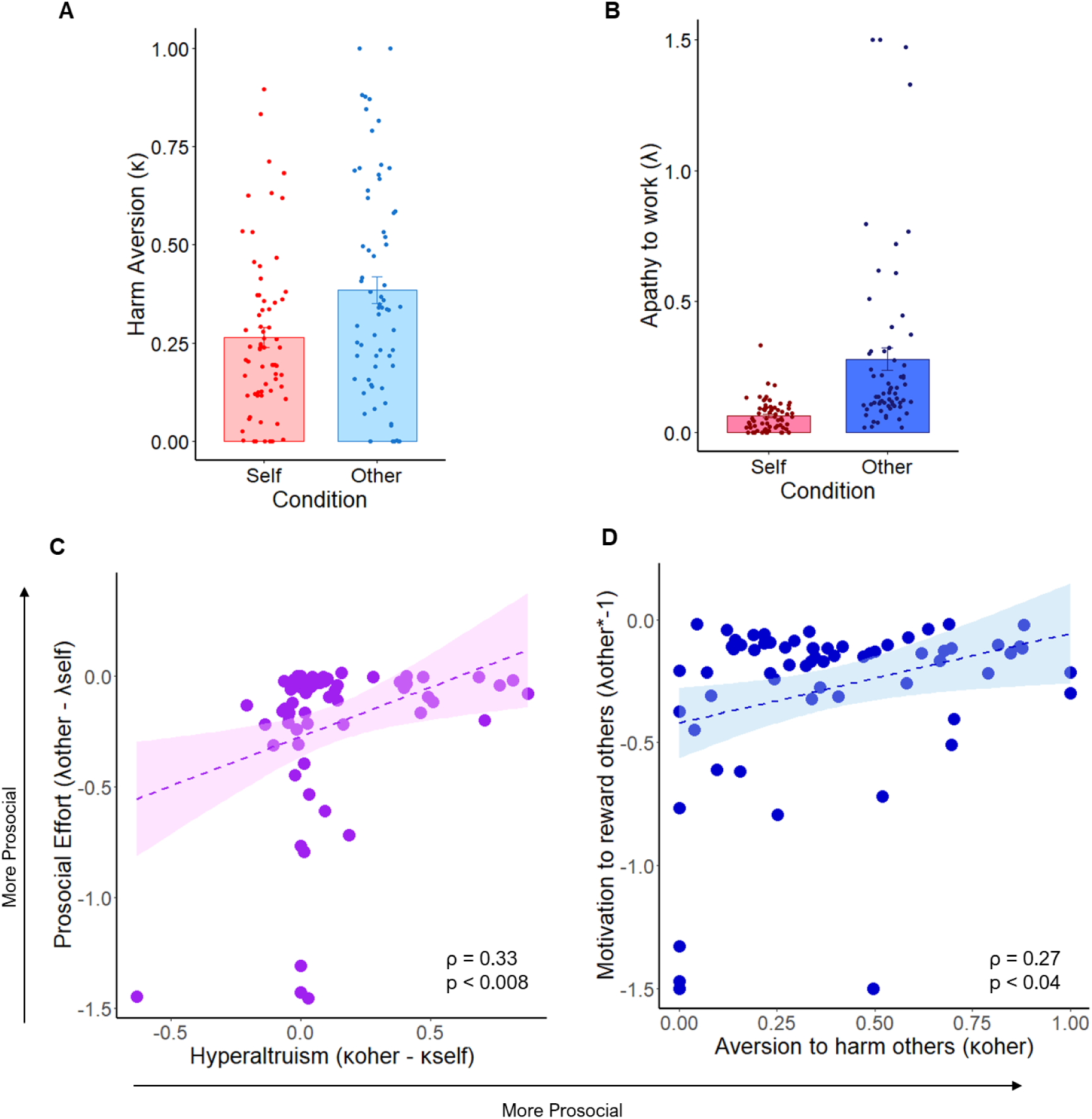
Prosocial behaviours across dimensions are correlated. A. Participants are more averse to harm others compared with themselves as measured by computational parameters (κ_other_ > κ_self_, p < 0.001). Y-axis depicts the value of the harm aversion parameter κ, which represents the degree in which profit is discounted by harm. B. Participants are more apathetic to exert effort for others compared with themselves as measured by computational parameters (λ_other_ > λ_self_, p < 0.001). Y-axis depicts the value of the discount parameter λ, which represents the degree in which reward is discounted by effort. C. Individual differences in prosocial preferences across tasks were associated: participants who were more averse to harming others relative to themselves in the harm aversion task (κ_other_ - κ_self_, x-axis) were also more willing to exert effort to benefit others relative to themselves in the prosocial effort task (λ_other_ - λ_self_, y-axis). D. Decisions to help others were positively correlated across tasks: participants who were more willing to exert effort to reward others (inversed λ_other_, y-axis) were also more averse to harming others (κ_other_, x-axis). Shaded areas indicate 95% confidence intervals. Dots represent individual participants.

Next, we tested whether prosocial behaviours across the two tasks were related— specifically, whether individuals who were more prosocial in one task also tended to be more prosocial in the other—replicating previous findings^6^. To do this, we reverse-coded the prosocial apathy effect (λ_other_ – λ_self_) and called this variable *prosocial effort*, such that positive values for both prosocial effort and hyperaltruism (κ_other_ – κ_self_) consistently reflected stronger prosocial preferences. A Spearman rank correlation revealed a significant association between prosocial effort and hyperaltruism (ρ = 0.33, p < 0.008, **Fig. 3C**), indicating that individuals who were more willing to exert effort for others were also more averse to harming them, relative to themselves. We then tested whether this relationship was driven specifically by preferences in other or self trials. To this end, we conducted Spearman partial correlations between κ_other_ and reversed-coded λ_other_ (controlling for their self counterparts), and between κ_self_ and reversed-coded λ_self_ (controlling for their other counterparts). Behavioural alignment was found only in other trials (ρ =0.27, p < 0.04, **Fig. 3D**) with no association in self trials (ρ = 0.02, p = 0.89, **Supplementary Fig. S2**). Finally, Contreras-Huerta et al.^6^ showed that the shared variance between hyperaltruism and prosocial effort was linked to a constellation of traits associated with emotional reactivity—specifically, negatively with emotional apathy and positively with affective empathy. We therefore correlated both prosocial indices with three apathy subscales^86^ and with cognitive and affective empathy^87^. We found that affective empathy was positively associated with hyperaltruism (ρ = 0.28, p < 0.03, **Supplementary Fig. S3A**), while emotional apathy was negatively associated with prosocial effort (ρ = -0.25, p < 0.05, **Supplementary Fig. S3B**). These results replicate and extend previous findings linking dispositional emotional sensitivity to domain-general prosocial preferences.

### Neural sensitivity to moral outcomes and their link to prosocial effort

Can neural sensitivity to moral outcomes explain individual differences in prosocial effort? To address this, we examined whether neural responses in the harm aversion task could be linked to the willingness to exert effort for others in a different context. Specifically, we tested whether individual differences in prosocial effort were predicted by neural sensitivity to anticipated harm to others or to profit gained from harming others. We focused on other trials from the harm aversion task, where decisions affected another person. A GLM was used to identify BOLD responses that varied parametrically with objective differences in money (Δm) and shocks (Δs) between harmful and helpful options. For control purposes, we also extracted statistical maps for Δm and Δs in self trials. Analyses targeted five a priori ROIs within the ACC, AI, and TPJ. To index prosocial motivation behaviourally, we used a mixed-effects model on other trials from the prosocial effort task, where decisions to work were predicted by reward, effort, and their interaction (see **equation 7**; for results of these models see **Supplementary Tables S3 and S4**). From this model, we extracted participant-level random slopes for reward—an established driver of prosocial motivation^54^. We then tested whether neural responses to Δm and Δs in the harm aversion task predicted reward sensitivity for others in the prosocial effort task, controlling for self-related slopes. This allowed us to evaluate two competing hypotheses: an other-regarding hypothesis, where greater neural sensitivity to others’ anticipated harm predicts increased reward sensitivity in the prosocial effort task, and a moral-conflict hypothesis, where greater neural sensitivity to profit gained by harming others is associated with lower reward sensitivity and reduced prosocial motivation.

First, we tested whether neural responses to others’ anticipated harm were associated with behavioural sensitivity to others’ rewards, as predicted by the other-regarding hypothesis. Specifically, we computed Spearman partial correlations between neural sensitivity to others’ harm (Δs_other_, controlling for Δs_self_) and the difference in reward sensitivity for others versus self, such that more positive values indicated greater parity in how participants valued others’ and self rewards (i.e., more prosocial motivation). All p-values were FDR-corrected across the five ROIs. Supporting the other-regarding hypothesis, we found a significant positive correlation in the vAI (**Fig. 4A**): greater neural responses to others’ harm were associated with higher relative valuation of others’ rewards (ρ = 0.36, p < 0.02 FDR-corrected; p = 0.003 uncorrected; **Fig. 4B**). This suggests that greater responses in the vAI to others’ harm was linked to more motivation to help others due to rewards. No other ROI showed significant associations, at either corrected or uncorrected thresholds. We next repeated the analysis using reward sensitivity for others only, controlling for reward sensitivity for self. Although this analysis did not survive FDR correction, the vAI showed a positive association with others’ rewards sensitivity with similar effect size (ρ = 0.30, p = 0.07 FDR-corrected; p < 0.02 uncorrected; **Fig. 4C**). To test robustness, we re-ran both analyses excluding participants with neural values ±3 SD from the mean. Both effects held and were strengthened under this exclusion criterion (ρ = 0.43, p = 0.002 FDR-corrected for the between-condition comparison; ρ = 0.41, p = 0.004 FDR-corrected for the other-only model; both p < 0.001 uncorrected). Together, these findings suggest that individuals with stronger vAI responses to others’ anticipated harm are more behaviourally responsive to others’ rewards—particularly when compared to self—supporting the notion of reduced prosocial apathy.

**Figure 4.**
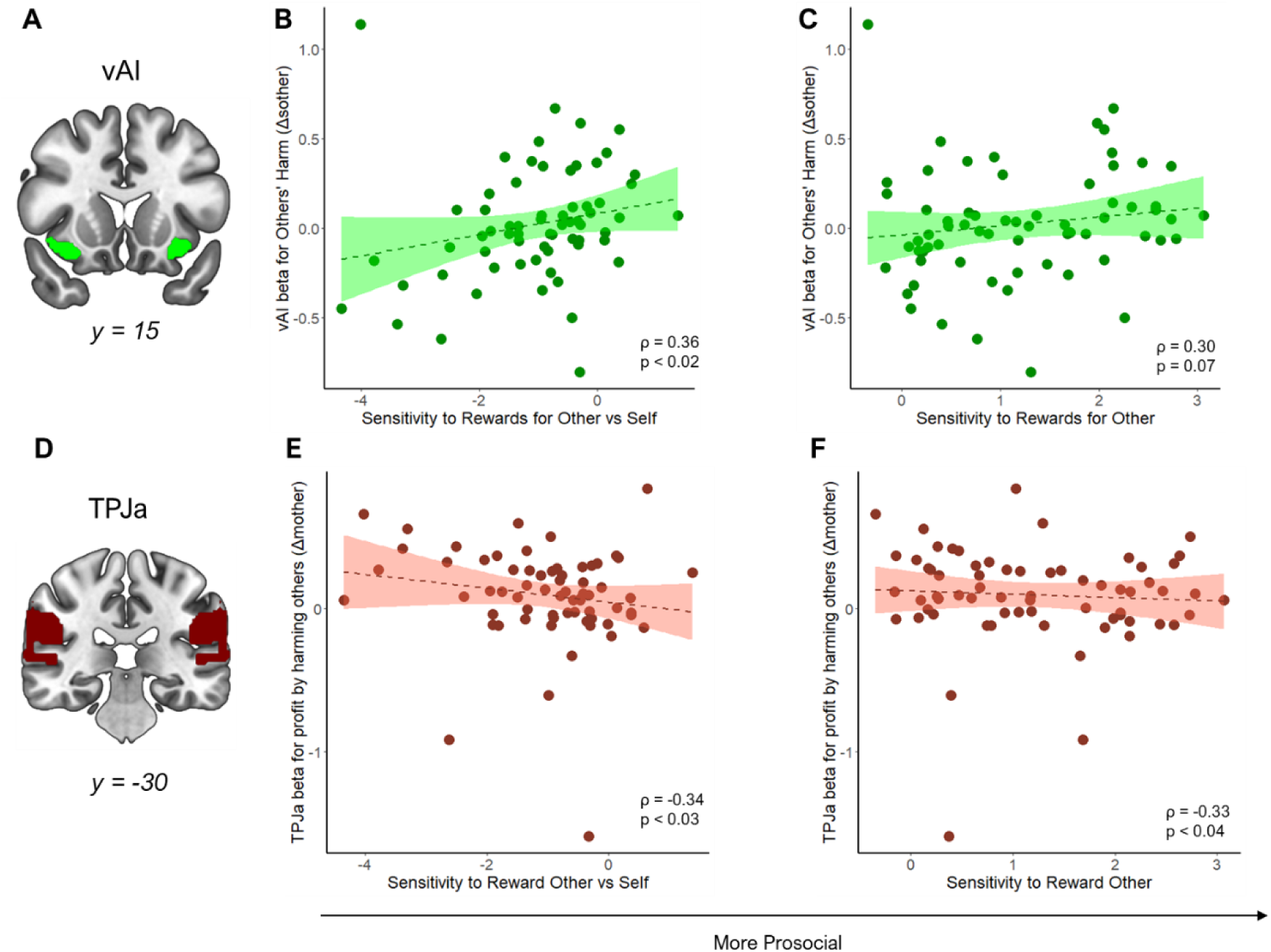
Neural responses to others’ harm and ill-gotten profit in the harm aversion task predict individual differences in reward sensitivity for others in the prosocial effort task. A. Mask of the ventral anterior insula (vAI), used to extract statistical maps for neural responses to anticipated harm (Δs_other_) and profit (Δm_other_) from harming others. B vAI responses to others’ harm (y-axis) were positively associated with behavioural reward sensitivity for others relative to self (x-axis), indicating stronger prosocial motivation. Reward sensitivity was indexed by participant-level beta slopes from mixed-effects models predicting work decisions in the prosocial effort task. C. vAI responses to others’ harm also showed a positive (uncorrected) association with reward sensitivity for others alone (controlling for self). D. Mask of the anterior temporo-parietal junction (TPJa), used to extract statistical maps for Δs_other_ and Δm_other_. E. TPJa responses to profit gained from harming others were negatively associated with reward sensitivity for others versus self— indicating reduced prosocial motivation and a tendency to prioritise self-benefit. F. TPJa responses to profit were also negatively associated with reward sensitivity for others (controlling for self). All p-values are FDR-corrected. Shaded areas represent 95% confidence intervals. Dots represent individual participants.

Next, we examined whether neural responses to profit gained by harming others were associated with behavioural sensitivity to others’ rewards, as predicted by the moral conflict hypothesis. Specifically, we tested Spearman partial correlations between Δm_other_ (controlling for Δm_self_) and the difference in reward sensitivity for others versus self. Consistent with the moral conflict hypothesis, neural responses in TPJa to profit (**Fig. 4D**) from harming others were significantly associated with lower reward sensitivity for others relative to self (ρ = -0.34, p < 0.03 FDR-corrected; p = 0.005 uncorrected; **Fig. 4E**). This suggests that individuals who show stronger neural responses to morally tainted gains in the TPJa tend to prioritise their own rewards over those of others. We then conducted a follow-up analysis correlating Δm_other_ with reward sensitivity for others only, controlling for reward sensitivity for self. This revealed a significant negative association in the same region: higher TPJa responses to ill-gotten profit predicted lower reward sensitivity for others (ρ = –0.33, p < 0.04 FDR-corrected; p = 0.007 uncorrected; **Fig. 4F**). To test the robustness of these results, we excluded participants whose neural responses were ±3 standard deviations from the mean. Both effects remained significant after this exclusion (ρ = -0.33, p < 0.04 FDR-corrected; p = 0.008 uncorrected for the first model; ρ = –0.34, p < 0.04 FDR-corrected; p = 0.007 uncorrected for the second model). Together, these findings tie the moral-conflict hypothesis to the TPJa: individuals who show stronger neural responses in this region to profit gained by harming others are less motivated to work for others’ benefit—highlighting a potential neural marker of reduced prosociality.

For completeness, we conducted a series of additional analyses to further support our findings. First, to explore alternative explanations, we tested whether Δm_other_ or Δs_other_ signals were correlated with individual differences in sensitivity to effort when obtaining rewards for others. We performed Spearman partial correlations using the same approach as our main analyses. None of these analyses revealed significant results at either corrected or uncorrected levels. Thus, alternative hypotheses—such as a valence-related account (i.e., responses to harm [Δs_other_] predicting effort sensitivity for others) and a self-centred account (i.e., responses to ill-gotten self profit [Δm_other_] predicting self-effort sensitivity)—were not supported. These findings suggest that harm aversion-related neural signals are more specifically associated with sensitivity to others’ benefits, rather than with effort sensitivity in prosocial contexts.

Second, although the focus of this study is on individual differences, we also conducted group-level analyses to test whether average responses to Δs_other_ in the vAI and Δm_other_ in the TPJa were significantly different from zero. These analyses revealed that responses to Δm_other_ in TPJa were significantly above zero (Wilcoxon rank-sum test, W = 1659, p < 0.001, M = 0.02, SEM = 0.04), whereas responses to Δs_other_ in vAI were not (W = 1157, p = 0.75, M = 0.10, SEM = 0.04). To complement this, we also performed a whole-brain analysis to identify clusters that significantly tracked harm and profit for self and other. These uncorrected results (p < 0.001, k > 30) are reported in **Supplementary Tables S5-S8.**

Finally, we replicated our neural-behaviour correlations using behavioural indices from the harm aversion task. We constructed separate mixed-effects models for self and other trials, predicting helping decisions from differences in money, shocks, and their interaction. From these, we extracted participant-level slopes for money and shocks, indexing sensitivity to ill-gotten profit and others’ harm. Fixed-effects results are reported in **Supplementary Tables S9 and S10**. We then tested whether these behavioural indices predicted reward sensitivity for others in the prosocial effort task, controlling for self-related slopes. Sensitivity to others’ harm was positively associated with reward sensitivity for others (ρ = 0.25, p < 0.05; **Supplementary Fig. S4A**), and sensitivity to profit from harming others also showed a significant positive correlation (ρ = 0.29, p =0.02; **Supplementary Fig. S4B**). Notably, this positive association reflects reduced temptation to harm for profit, as money differences negatively influenced helping decisions (more positive slopes indicate stronger resistance to ill-gotten gains). Together, these behavioural results align with and reinforce our neural findings. For completeness, we also tested correlations between effort sensitivity and harm/profit sensitivity, but these analyses yielded no significant effects.

Taken together, these results suggest that neural signals associated with moral harm aversion in specific regions are selectively linked to how individuals value others’ outcomes in a different domain of prosocial behaviour. Stronger vAI responses to anticipated harm were associated with greater valuation of others’ rewards, whereas TPJa responses to profit from harming others were associated with reduced valuation of others’ welfare. Crucially, these associations were not explained by general sensitivity to effort, indicating that harm aversion reflects how people differentially value others’ welfare beyond the physical costs of acting prosocially.

## DISCUSSION

Prosocial behaviour can take multiple forms, from avoiding harm to others to exerting effort to secure their rewards. Although people are generally more motivated to prevent harm than to actively benefit others^4,5,18,19,23,24^, it remains unclear whether sensitivities to moral outcomes in one context can predict prosocial tendencies in another. In this study, we show that neural and behavioural responses to others’ harm and to ill-gotten profits are linked to motivation to obtain rewards for others in a separate domain—albeit in opposite directions. Greater vAI responses to anticipated harm predicted stronger motivation to obtain others’ rewards, whereas stronger TPJa responses to profit from harming others predicted reduced sensitivity to work for rewards for others. These findings suggest that distinct neural mechanisms—one associated with sensitivity to others’ suffering and another with processing moral conflict—can shape prosocial behaviour across different domains.

Our finding that vAI responses to others’ suffering predicted greater motivation to work for others’ rewards suggests that this region may act as a socio-affective hub that links moral sensitivity with prosocial motivation across contexts^31,40,42,44,49,50,55,56^ . Importantly, this effect was specific to the vAI and not observed in the dAI, highlighting functional differences within the insula. Whereas the dAI is structurally connected to executive networks (including the medial and dorsomedial prefrontal cortices and superior parietal regions) and implicated in attention and cognitive control, the vAI is more closely connected to limbic regions and the ACCg, supporting its role in affective and social processing^78,79,81,88–90^. Consistent with this, the vAI has been proposed to encode empathic representations of others’ states, and heightened vAI responses have been linked to prosocial tendencies, including extraordinary prosociality^41,91^. By contrast, the dAI may be less directly involved in evaluating social outcomes relevant to prosocial motivation and more engaged in self–other distinction processes, in line with its role in bodily awareness and interoception^54,92^. Taken together, these findings suggest that the vAI supports prosocial decision-making by representing others’ affective outcomes, providing a mechanism through which sensitivity to others’ harm can generalise into motivation to obtain their rewards.

Further highlighting subregional specialisation in prosocial behaviour, TPJa appears to play a distinctive role in signalling moral conflict during prosocial decision-making. In our data, lower TPJa responses to ill-gotten rewards—profits earned by harming others—were associated with stronger motivation to seek others’ rewards a week later. In contrast with the more other-regarding involvement of the vAI, TPJa may reflect the tension between self- and other-interests. Individuals whose TPJa responses to ill-gotten rewards were less attenuated appeared less inclined to work for others’ rewards. This finding contributes to ongoing debates about TPJ function in social behaviour: some studies link TPJ activity to other-regarding motives^49–51,69^, whereas others highlight its role in signalling conflict during morally costly decisions. Our results suggest that TPJ may be sensitive to social and moral information that can modulate reward-related processing^19,24,65,68,93^, with this function being particularly associated with its anterior subregion (TPJa). The interpretation of TPJa as s primarily supporting a moral conflict role aligns with accounts proposing a functional subdivision within the TPJ. TPJa has been linked to attentional reorienting and cognitive control processes^94,95^, functions that may be critical for detecting moral norm violations or evaluating conflicts between competing motives. In contrast, its posterior portion (TPJp) has been more strongly implicated in theory of mind and cognitive empathy—processes central to understanding others’ beliefs, intentions, and perspectives ^60,95^. Thus, whereas TPJp may support prosociality by enabling mental state attribution and perspective-taking, TPJa may influence prosocial decisions by signalling the moral tension inherent in choosing between self-benefit and harm to others.

Even though harm aversion and prosocial effort were correlated, our behavioural results suggest that they are driven by partially distinct motivational processes at the task level. In the harm-aversion task, participants were more reluctant to profit from harming others than from harming themselves. This pattern was partly explained by greater profit sensitivity when the costs were personal, since monetary gains were more influential when the shocks were self-directed than when they harmed others—consistent with a motivation to avoid morally costly actions^19^. By contrast, in the prosocial-effort task, participants were more willing to exert effort for their own rewards than for others’, reflecting the joint influence of reward and effort sensitivity. Because prosocial effort lacks an explicit moral imperative (such as “do no harm”), prosocial behaviour in this context depends on the extent to which individuals value others’ outcomes relative to their own effort costs.

Importantly, despite these task-level differences, individual variability in neural and behavioural responses to others’ harm predicted greater subsequent prosocial effort. This convergence suggests that trait-like sensitivity to others’ welfare, as reflected in vAI activity, may underlie a general disposition toward prosociality. Although vAI activity is not always directly linked to harm-aversion behaviour^19^, its variability across individuals explained prosocial effort, supporting the idea that people differ in how much they value others’ outcomes across contexts. This interpretation aligns with evidence that empathic traits modulate neural sensitivity to others’ benefits in prosocial behaviour^34,40,41,96,97^, as well as with previous findings linking harm aversion and prosocial effort through shared emotional reactivity^6^.

Extending beyond the vAI, our results show that TPJa activity in response to moral conflict also predicts behaviour in a task without an explicit moral frame (i.e., prosocial effort). This suggests that moral principles may generalise accross domains, influencing behaviour even when moral norms are not directly at stake. Philosophical and empirical accounts alike have proposed that moral identity and self-consistency guide actions in different contexts^98–100^, and individuals often strive to reduce discrepancies between their moral standards and behaviour to avoid guilt^73,101^. Indeed, we recently found that larger gaps between people’s moral beliefs (as judged in third-person scenarios) and their own behaviour are linked to reduced prosocial motivation^74^ . Thus, individual differences in conflict-related responses within TPJa may reflect stable dispositions that shape broader prosocial tendencies. This may be supported by affective mechanisms, given that immoral actions reliably elicit strong negative emotions such as disgust^102–104^—consistent with affective reactivity underpinning both prosocial effort and harm aversion^6^. One limitation of the present study is that the link between harm aversion and prosocial effort was inferred from cross-task correlations measured a week apart, rather than from neural signals recorded during prosocial-effort decisions themselves. Future work should examine how neural signals relate across tasks and whether shared mechanisms underlie them. TPJ and AI have been shown to track subjective value of working for both self and other’s rewards^24^, and it would be interesting to test whether these areas constitute the core of a common prosocial disposition. It will be also important to test how neural sensitivity to others’ benefits interacts with self-costs when both carry negative valence—for example, when people exert effort to prevent harm or endure harm to help others. Recent evidence shows that the typical “prosocial apathy” effect diminishes when effort serves to avoid causing harm^105^, highlighting the influence of moral framing. Another open question is whether these moral neural sensitivities generalise to predict prosociality across diverse domains, including charitable giving, cooperation in economic games, or spontaneous helping in naturalistic settings^28,48,106^. Crucially, it remains unclear whether individuals who show strong neural moral signatures in laboratory tasks also display consistent prosocial behaviour in daily life. Addressing this gap will require ecologically valid paradigms and multi-method approaches, including experience sampling and digital phenotyping. These efforts are not only theoretically important but also socially relevant. A clearer understanding of the motivational and neural foundations of prosociality could inform interventions to foster cooperative and compassionate behaviour —for example, by leveraging affective engagement and moral consistency to promote prosociality in everyday contexts^1,107–109^.

In summary, we found that individual differences in vAI responses to others’ harm were associated with greater sensitivity to others’ rewards in effort-based decisions, suggesting that this region tracks morally salient outcomes that influence prosocial motivation across contexts. In contrast, TPJa responses to ill-gotten gains were linked to reduced prosocial effort, consistent with a role in detecting moral conflict between self- and other-interest. Together, these findings indicate that neural sensitivity to morally salient information can predict prosocial behaviour in a distinct domain, and highlight the importance of individual variability in shaping moral and prosocial tendencies.

## DATA AND CODE ACCESSIBILITY

All data and scripts used for main analysis and figures can be found here https://osf.io/a3rey/?view_only=09cb6bc7f6d04a6e9eb5a7051b04cdc3.

## AUTHORS’ CONTRIBUTIONS

LSCH: Conceptualization, Methodology, Software, Validation, Formal analysis, Investigation, Data Curation, Writing - Original Draft, Writing - Review & Editing, Visualization, Project administration. HY: Methodology, Investigation, Software, Data curation, Writing – review & editing. AMBP: Methodology, Investigation, Writing – review & editing. PLL: Methodology, Writing - Review & Editing. FRT: Writing – review & editing. MJC: Conceptualization, Methodology, Resources, Writing – review & editing, Supervision, Project administration, Funding acquisition. MAJA: Conceptualization, Methodology, Writing – original draft, Writing – review & editing, Supervision.

