## Supplementary Information for "Neural signatures of harm aversion predict later willingness to exert effort for others’ rewards"

### SUPPLEMENTARY METHODS: SOCIAL MANIPULATION PROCEDURE

Participants completed two types of trials in both the fMRI and prosocial effort tasks: self trials, where they were the beneficiary of the reward, and other trials, where an unknown person was the beneficiary. To ensure that participants believed their decisions had real consequences for others, we implemented a rigorous social manipulation protocol, previously validated to establish trust in the authenticity of social trials (1, 2).

**Social Manipulation Protocol (fMRI Session).** The social manipulation was framed as a role assignment procedure. Before entering the scanner, participants were informed that two people were participating in the study, each assigned one of two roles: Decider or Receiver. The Decider would perform decision-making tasks affecting both themselves and the Receiver, while the Receiver would complete an unrelated set of tasks with no personal benefit. Participants were told that roles would be assigned randomly by drawing a ball from a box. However, unbeknownst to them, all participants were assigned the role of Decider, while the Receiver was a confederate. After data collection, all participants were debriefed via email about this deception.

The protocol followed a structured sequence: (i) *Preparation*: The experimenter explained the role assignment procedure in the experimental room. (ii) *Random Assignment*: A second experimenter arrived with the confederate, standing on opposite sides of a semi-opened door to prevent visual identification. Both individuals were instructed not to speak and were handed a yellow rubber glove to obscure identifying features. (iii) *Mutual Awareness*: Participants and the confederate waved their gloved hands at each other through the door to confirm the presence of another person. (iv) *Role Assignment*: Participants and the confederate drew a ball from the box, with the selection order determined by a coin toss. The second experimenter and the confederate then exited, and participants were informed that they had been assigned the Decider role.

To minimise reputation concerns, social desirability bias, and reciprocity effects—all of which could influence social behaviour (3, 4)—participants were explicitly told that: their choices were anonymous, they would never meet or interact with the Receiver, and Receivers would be performing a separate, unrelated task in parallel. To assess belief in the social manipulation, participants answered debriefing questions at the end of each session. These subtle questions were designed to gauge their trust in the procedure without directly prompting doubt about its validity.

**Social Manipulation Protocol (Behavioural Session).** A second social manipulation was introduced in the behavioural session of the experiment. At the beginning of the prosocial effort task, participants received instructions and completed the calibration procedure (see Materials and Methods in the main text). Next, they were told that they would be paired with a Receiver from the previous fMRI session, though not necessarily the same one. A second experimenter then entered the room, and, as in the fMRI session, both the participant and the confederate stood on opposite sides of a semi-opened door, waving their hands to establish mutual awareness. After this interaction, the second experimenter and the Receiver left, and participants proceeded with the task.

**SUPPLEMENTARY TABLES**

**Table S1.** Results of the harm aversion model predicting trial-by-trial helping decisions. The model included the difference in money, the difference in shocks, the recipient of the shocks, and their interactions as predictors (see Materials and Methods for details).

| *Effect* | *beta* | *SEM* | *z* | *p* |
|---|---|---|---|---|
| *Δmoney* | -4.22 | 0.40 | -10.67 | < 0.001 |
| *Δshocks* | 1.61 | 0.18 | 9.17 | < 0.001 |
| *Recipient* | 1.32 | 0.09 | 15.44 | < 0.001 |
| *Δmoney * Recipient* | 0.62 | 0.09 | 6.86 | < 0.001 |
| *Δshocks * Recipient* | 0.12 | 0.10 | 1.45 | 0.15 |
| *Δmoney * Δshocks* | < 0.00 | 0.07 | -0.01 | 0.99 |
| *Δmoney * Δshocks * Recipient* | 0.12 | 0.09 | 1.34 | 0.18 |

**Table S2.** Results of the prosocial effort model predicting trial-by-trial decisions to work or rest. The model included effort level, reward magnitude, beneficiary of the reward, and their interactions as predictors (see Materials and Methods for details).

| *Effect* | *beta* | *SEM* | *z* | *p* |
|---|---|---|---|---|
| *Reward* | 2.22 | 0.18 | 12.08 | < 0.001 |
| *Beneficiary* | -3.45 | 0.11 | -32.13 | < 0.001 |
| *Effort* | -2.52 | 0.21 | -11.82 | < 0.001 |
| *Reward*Beneficiary* | -0.93 | 0.10 | -9.62 | < 0.001 |
| *Reward*Effort* | -0.01 | 0.08 | -0.14 | 0.89 |
| *Beneficiary*Effort* | 0.26 | 0.10 | 2.61 | 0.009 |
| *Reward*Beneficiary*Effort* | -0.03 | 0.09 | -0.35 | 0.73 |

**Tables S3 y S4.** Results of the prosocial effort models predicting trial-by-trial helping decisions separately for self and other. The model included effort level, reward magnitude, and their interaction as predictors (see Materials and Methods for details).

***Table S3. Self Model***

| *Effect* | *beta* | *SEM* | *z* | *p* |
|---|---|---|---|---|
| *Reward* | 2.38 | 0.25 | 9.60 | < 0.001 |
| *Effort* | -2.51 | 0.25 | -10.26 | < 0.001 |
| *Reward*Effort* | 0.15 | 0.11 | 1.36 | 0.18 |

***Table S4. Other Model***

| *Effect* | *beta* | *SEM* | *z* | *p* |
|---|---|---|---|---|
| *Reward* | 1.25 | 0.15 | 8.20 | < 0.001 |
| *Effort* | -2.42 | 0.18 | -13.87 | < 0.001 |
| *Reward*Effort* | -0.15 | 0.07 | -2.26 | 0.02 |

**Tables S5 – S8.** Uncorrected (p < 0.001, k > 30) whole brain results for differences in money and shocks between the harmful and the helpful options for *self* and *other* trials in the harm aversion task.

***Table S5. Differences in shocks in self trials***

| *Regions* | *x* | *y* | *z* | *F* | *k* |
|---|---|---|---|---|---|
| *Lingual Gyrus* | 12 | -79 | -7 | 79.99 | 343 |
| *Superior Frontal Gyrus* | -18 | -10 | 65 | 39.1 | 1107 |
| *Precuneus* | 9 | -67 | 41 | 25.34 | 144 |
| *Central Opercular Cortex* | -54 | -1 | 5 | 24.4 | 167 |
| *Parietal Operculum Cortex* | 57 | -31 | 26 | 24.07 | 102 |
| *Precentral Gyrus* | 63 | 8 | 38 | 23.73 | 94 |
| *Supramarginal Gyrus* | -48 | -19 | 26 | 21.15 | 101 |
| *Inferior Frontal Gyrus* | -30 | 35 | 17 | 25.84 | 61 |
| *Right Putamen* | 27 | -1 | 14 | 25.67 | 72 |
| *Supramarginal Gyrus* | 66 | -34 | 50 | 23.98 | 37 |
| *Lingual Gyrus* | 0 | -70 | 5 | 22.84 | 43 |
| *Superior Temporal Gyrus* | 72 | -28 | 17 | 20.43 | 46 |
| *Cerebellum VIIIa* | -18 | -67 | -49 | 19.33 | 58 |
| *Lateral Occipital Cortex* | 39 | -82 | 8 | 18.86 | 39 |
| *Precentral Gyrus* | 48 | -4 | 59 | 18.14 | 33 |

***Table S6. Differences in money in self trials***

| *Regions* | *x* | *y* | *z* | *F* | *k* |
|---|---|---|---|---|---|
| *Angular Gyrus* | 51 | -52 | 20 | 33.88 | 474 |
| *Angular Gyrus* | -45 | -55 | 20 | 23.66 | 312 |
| *Superior Temporal Gyrus* | 57 | -1 | -16 | 22.93 | 150 |
| *Superior Temporal Gyrus* | -51 | -31 | 2 | 21.7 | 93 |
| *Right Amygdala* | 24 | -1 | -19 | 21.83 | 53 |
| *Left Amygdala* | -18 | -7 | -19 | 20.12 | 42 |
| *Posterior Cingulate Gyrus* | -9 | -43 | 38 | 15.62 | 72 |

***Table S7. Differences in shocks in other trials***

| *Regions* | *x* | *y* | *z* | *F* | *k* |
|---|---|---|---|---|---|
| *Lingual Gyrus* | -12 | -82 | -13 | 48.27 | 450 |
| *Left Caudate* | -15 | -7 | 26 | 19.78 | 38 |
| *Superior Parietal Lobule* | 42 | -43 | 68 | 19.24 | 117 |
| *Right Caudate* | 21 | -4 | 26 | 19.15 | 35 |
| *Precentral Gyrus* | 54 | 8 | 23 | 18.9 | 38 |
| *Cerebellum VIIb* | -30 | -73 | -58 | 17.69 | 35 |
| *Postcentral Gyrus* | -45 | -37 | 65 | 15.29 | 37 |
| *Precentral Gyrus* | 33 | -7 | 65 | 14.34 | 35 |

***Table S8. Differences in money in other trials***

| *Regions* | *x* | *y* | *z* | *F* | *k* |
|---|---|---|---|---|---|
| *Lateral Occipital Cortex* | -54 | -67 | 26 | 33.45 | 273 |
| *Superior Parietal Lobule* | 24 | -55 | 50 | 23.26 | 39 |
| *Frontal Pole* | -12 | 59 | 20 | 23.25 | 42 |
| *Superior Frontal Gyrus* | -24 | 35 | 44 | 18.55 | 34 |
| *Angular Gyrus* | 42 | -49 | -29 | 18.22 | 62 |

**Tables S9 y S10.** Results of the harm aversion models predicting trial-by-trial helping decisions separately for self and other. The model included the difference in money, the difference in shocks and their interactions as predictors (see Materials and Methods for details).

***Table S9. Self Model***

| *Effect* | *beta* | *SEM* | *z* | *p* |
|---|---|---|---|---|
| *Δmoney* | -5.38 | 0.55 | -9.81 | < 0.001 |
| *Δshocks* | 1.77 | 0.19 | 9.16 | < 0.001 |
| *Δmoney* Δshocks* | -0.04 | 0.10 | -0.37 | 0.72 |

***Table S10. Other Model***

| *Effect* | *beta* | *SEM* | *z* | *p* |
|---|---|---|---|---|
| *Δmoney* | -3.52 | 0.37 | -9.58 | < 0.001 |
| *Δshocks* | 1.86 | 0.19 | 10.09 | < 0.001 |
| *Δmoney* Δshocks* | 0.03 | 0.08 | -0.34 | 0.73 |

### SUPPLEMENTARY FIGURES

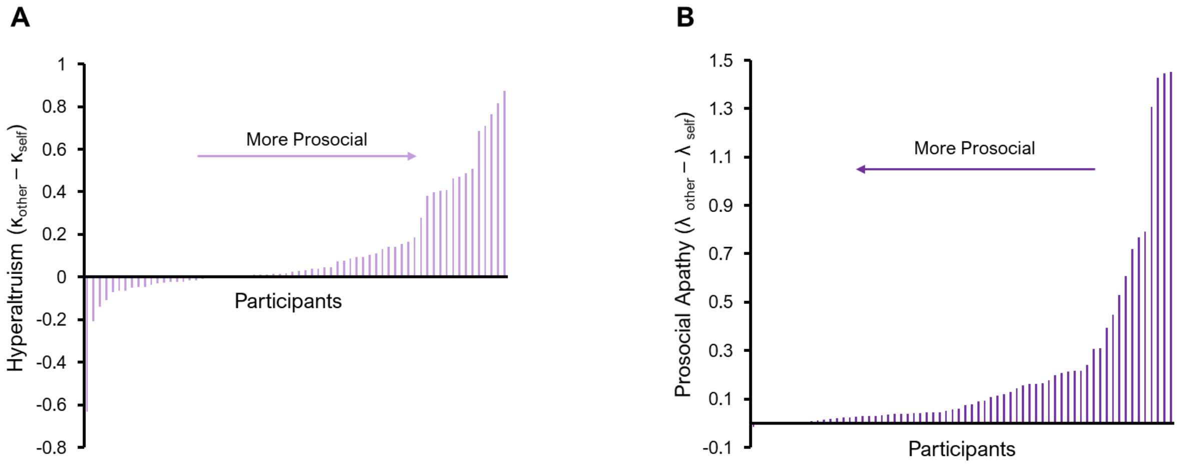


**Supplementary Figure S1. People show different behavioural trends in harm aversion and prosocial effort contexts.** A. The hyperaltruism effect ($\kappa_{other} > \kappa_{self}$, y-axis) is seen in 62.5% of the current sample. The x-axis represents each individual ordered from lowest to highest values of hyperaltruism. B. The prosocial apathy effect ($\lambda_{other} > \lambda_{self}$, y-axis) is seen in 95.5% of the current sample. The x-axis represents each individual ordered from lowest to highest values of prosocial apathy

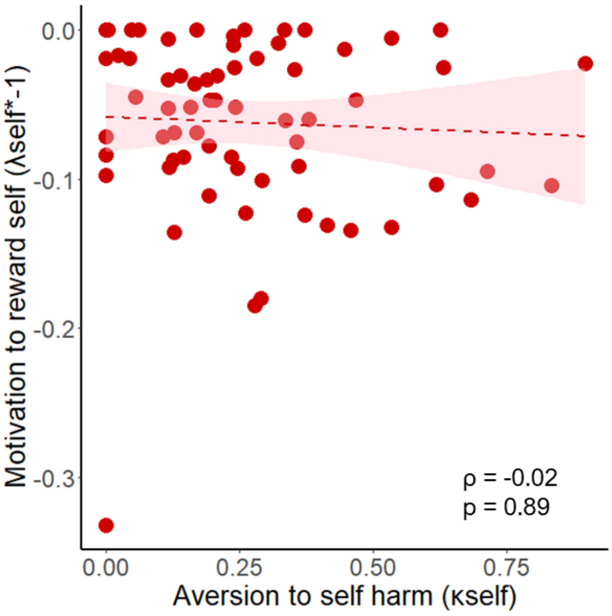


**Supplementary Figure S2. No significant association was observed between decisions to benefit the self across tasks.** Participants who were more willing to work for self-reward (reversed $\lambda_{self}$, y-axis) were not more averse to harming themselves ($\kappa_{self}$, x-axis). Shaded areas indicate 95% confidence intervals. Dots represent individual participants.

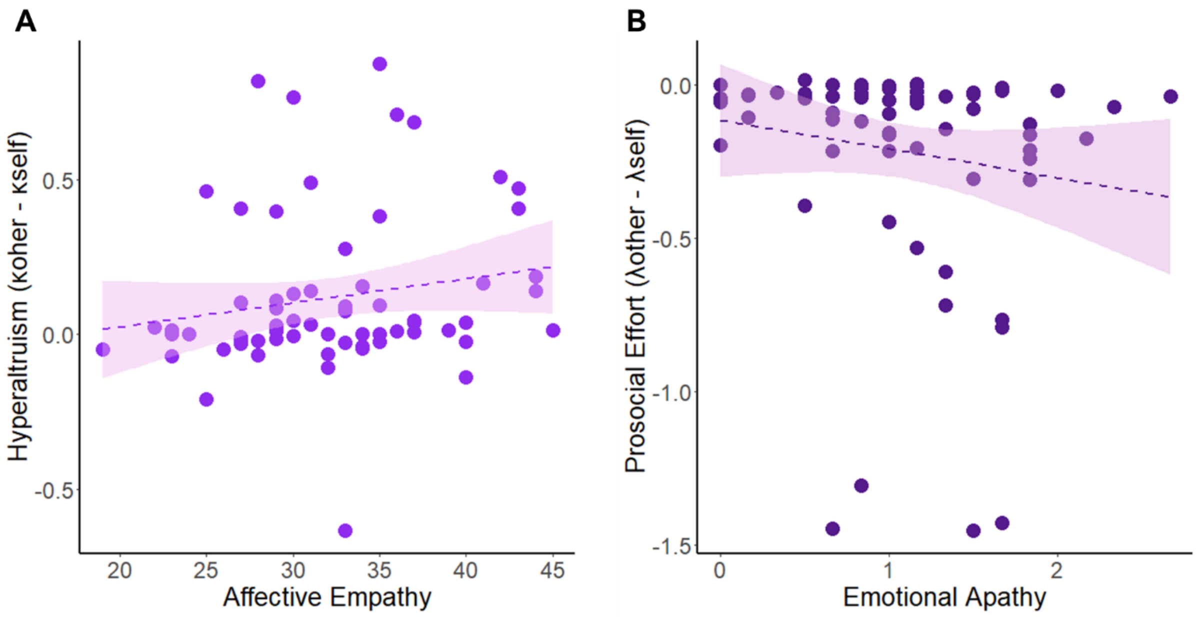


**Supplementary Figure S3. Associations between prosocial behaviours and affective traits.** A. Hyperaltruism in the harm aversion task is positively correlated with Affective Empathy, as measured by the Questionnaire of Cognitive and Affective Empathy (QCAE; Reniers et al., 2011), ρ = 0.28, p = 0.02. B. Prosocial effort in the effort task is negatively correlated with Emotional Apathy, as measured by the Apathy-Motivation Index (AMI; Ang et al., 2017), ρ = –0.25, p < 0.05.

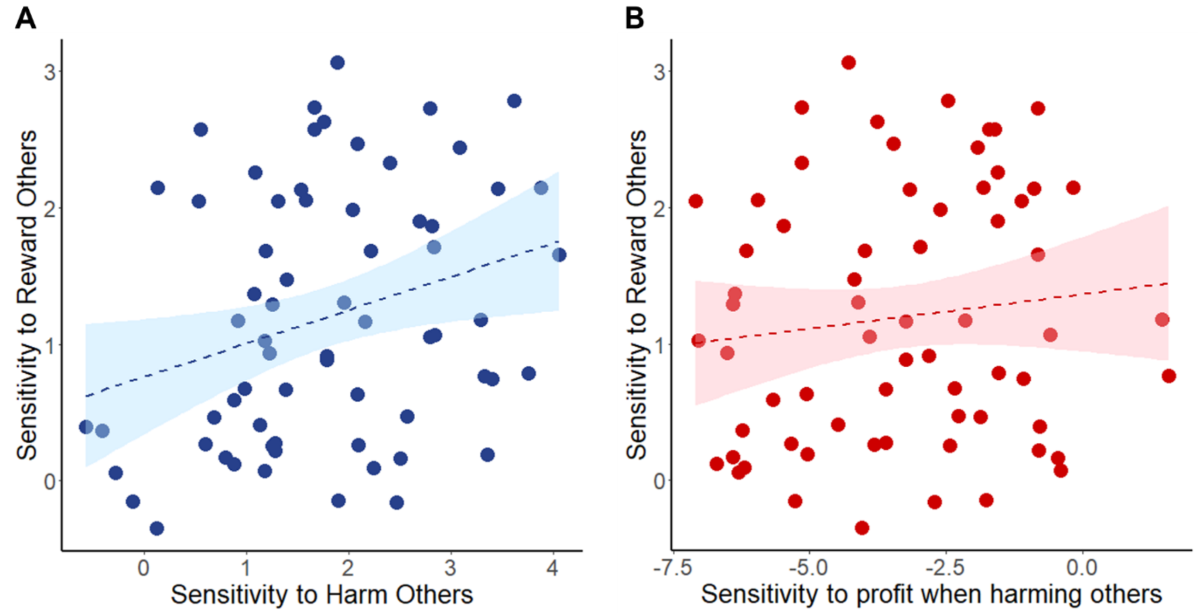


**Supplementary Figure S4. Behavioural sensitivity to harm and profit in the harm aversion task predicts reward sensitivity for others in the prosocial effort task.** A. Sensitivity to others' harm (x-axis)—indexed by individual slopes for shock differences in the harm aversion task—was positively associated with reward sensitivity for others (y-axis) in the prosocial effort task ($\rho = 0.25$, $p < 0.05$). B. Sensitivity to profit from harming others (x-axis)—indexed by individual slopes for money differences—was also positively correlated with reward sensitivity for others ($\rho = 0.29$, $p < 0.03$). Note: This positive association reflects greater resistance to ill-gotten gains, as higher money slopes indicate weaker influence of profit on helping decisions (i.e., reduced temptation to harm for monetary gain). Shaded areas represent 95% confidence intervals; each point represents an individual participant.
